# Effects of Simulated Microgravity on Human Hematological and Anemia-Related Biomarkers: A Systematic Review and Meta-Analysis

**DOI:** 10.1101/2025.09.17.676920

**Authors:** Ahmed Ayman Mohamed Abolyazed, Wafaa Ahmed Elbehery, Hanan Mahmoud Elsayed, Yousef Hamdy ElSheeta, Abdalla Yasser Elnemr

## Abstract

**Background:** Space anemia has been observed in astronauts after spaceflights for a long time, but its exact mechanism remains unclear. Because of the practical and ethical challenges of performing human studies in space, researchers use ground-based analogs to study the hematological changes that occur upon exposure to microgravity. Understanding these changes is important for protecting astronauts during long spaceflights and for guiding medical treatment on Earth in cases of prolonged inactivity or bed rest.

**Methods:** The literature search was done in April 2025 in PubMed, Web of Science, Scopus, and Cochrane. The inclusion criteria are randomized or non-randomized controlled trials and experimental repeated-measures studies that enrolled humans exposed to simulated microgravity for at least 24 hours. This study followed PRISMA (2020) guidelines and was registered in PROSPERO under CRD420251104161. Data was analyzed by Comprehensive Meta-Analysis software.

**Results:** Ten studies, involving 152 participants, met the inclusion criteria. During exposure, the first effect was a decreased plasma volume, leading to hemoconcentration. Erythropoietin concentration decreased, while parameters dependent on plasma volume, such as hemoglobin, hematocrit, and RBC count, falsely increased. Total hemoglobin mass and red cell volume showed a marked decrease, suggesting suppressed erythropoiesis. Iron status markers increased significantly, while classic hemolysis markers did not confirm increased hemolysis. Post-reambulation, erythropoietin rebounded, while the true decreases in hemoglobin, hematocrit, and RBC count were revealed as hemoconcentration resolved.

**Conclusion:** Simulated microgravity induces hemoconcentration, erythropoietin suppression, and altered iron metabolism. Further investigation is needed to clarify the role of hemolysis and to develop efficient countermeasures.

## 1. Introduction

Prolonged exposure to spaceflight causes multiple physiological changes in the human body. Microgravity, exposure to radiation, physical and emotional stress, nutritional changes, and disrupted circadian cycles affect many of the body’s functions, such as vision, the health of muscles and bones, and the immune system. (Williams et al. 2009). An important physiological change that occurs is fluid redistribution as a result of microgravity. This redistribution causes various changes in different hematological parameters, and these changes are referred to as space anemia.

The history of space anemia started in the 1960s following missions in NASA’s Gemini program when reductions in red blood cell mass were documented after returning to the earth (Fischer et al. 1967). These early findings were the core for recognition of space anemia as a physiological response to spaceflight.

Even though space anemia has been known about for a long time, the exact mechanism is still not fully understood. Different hypotheses were proposed to explain the reduction in red blood cell mass following exposure to microgravity. Suggested mechanisms include decreased erythropoietin (EPO) production, suppression of erythropoiesis, enhanced hemolysis, sequestration, and RBC dysfunction, but the precise physiological basis of space anemia remains unclear and not fully established (Trudel et al. 2022).

As humanity prepares for longer space missions, understanding the exact physiological responses of exposure to the space environment becomes essential, and among these responses, space anemia is considered a key concern (Lansiaux et al. 2024). This is crucial for both men and women as the number of female astronauts has progressively increased since the 1980s (Corlett et al. 2020). Deeper understanding of this condition is vital for developing suitable countermeasures to support the travelers during these prolonged missions.

Studying the effect of microgravity on different physiological aspects in space faced logistical, financial, and ethical challenges (Shelhamer et al. 2020), so to overcome these limitations, different ground-based models have been developed to simulate the effects of microgravity under controlled conditions. Among the most commonly used are head-down tilt bed rest and dry immersion, which not only mimic the physiological effects of microgravity but also offer a high degree of experimental control, which helps the researchers isolate specific variables and study them with greater precision (Watenpaugh 2016).

In addition to the great benefits in the field of space medicine, understanding the effect of models like head-down tilt bed rest and dry immersion on the hematological parameters can also have important clinical applications on earth. These models are used to mimic the effects of prolonged physical inactivity, which is a very common problem among various groups of patients, like the elderly and those with chronic immobilization, providing us with more knowledge about hematological and systemic changes associated with extended bed rest. Improved understanding of these changes can help us develop more strategies for preventing or managing anemia and support overall health in these vulnerable groups. (Strollo et al. 2018).

Despite the increasing number of studies about the effects of simulated microgravity on anemia-related biomarkers, the findings were inconsistent with variability in study designs, participant characteristics, and, most importantly, reported outcomes.

Previous reviews discussed the available literature, but this study quantitatively synthesizes the evidence, determines the overall effect size, and assesses changes in hematological biomarkers both during and after exposure to simulated microgravity, aiming to advance current understanding of microgravity-induced anemia and support the development of both space mission countermeasures and Earth-based clinical applications.

## 2. Methods

This study was conducted in accordance with the Preferred Reporting Items for Systematic Reviews and Meta-Analyses (PRISMA) 2020 guidelines (Page et al. 2021). We followed the Cochrane Handbook guidelines in doing all the steps (Higgins et al. 2019). The protocol was registered in the International Prospective Register of Systematic Reviews (PROSPERO) under the registration number CRD420251104161.

### 2.1 Inclusion criteria

Randomized or non-randomized controlled trials and experimental repeated-measures studies that enrolled human participants exposed to simulated microgravity for at least 24 hours were eligible. Included studies had to provide quantitative data on anemia-related hematological markers. Only full-text articles published in English were considered, with no restrictions on publication date, due to the limited number of studies available in this field.

### 2.2 Exclusion criteria

Studies that used animals, in vitro or computational models, didn’t have any microgravity exposure, were observational, case reports/series, conference abstracts, editorials, commentaries, or didn’t report extractable numerical data for the outcomes we were looking for were excluded.

### 2.3 Information Sources and Search Strategy

Two authors conducted a comprehensive literature search in four major electronic databases: PubMed, Cochrane Library, Scopus, and Web of Science. The search was performed in April 2025. No filters were applied during the search to ensure broad coverage. Reference lists of included studies were also screened manually to identify additional eligible records. The complete search strategy is provided in the supplementary material.

### 2.4 Study selection

The results of the literature search were collected in an Excel sheet and screened through the title and abstract by two independent authors. Then, the retrieved studies were further screened through full text to determine the final included studies. A third author was consulted to resolve any conflict.

### 2.5 Data extraction

Data extraction was performed by two reviewers separately and revised by a third reviewer. The reviewers collected general study information (study design, type of simulated microgravity, exposure duration, sample size, participant gender, presence or absence of a control group for microgravity or for other interventions, and the nature of such interventions if applicable). Baseline participant characteristics were also extracted, including age, body mass index (BMI), height, weight, and VO₂ max.

Regarding outcomes, the reviewers extracted quantitative data for hematological markers, which are hemoglobin, hematocrit, erythropoietin (EPO), plasma volume (PV), red cell volume (RCV), red blood cell (RBC) count, reticulocytes, total hemoglobin mass (tHb mass), mean corpuscular volume (MCV), mean corpuscular hemoglobin (MCH), mean corpuscular hemoglobin concentration (MCHC), serum iron, transferrin saturation, total and indirect bilirubin, and haptoglobin.

Each outcome was extracted at two primary time points: (1) baseline during the ambulatory period and (2) during or immediately after exposure to microgravity to assess the impact of the intervention. When available, a third post-recovery time point was also extracted, which is defined as 24 hours or more after reambulation to normal gravity. When multiple values were reported for the same outcome on different days within the same time window, we selected the most representative value of the response during that phase. The same approach was applied uniformly across all included studies.

### 2.6 Data Processing and Transformation

All outcome data were standardized to mean ± standard deviation (SD) before analysis. For the studies that reported standard errors (SE) (Dunn et al., 1984; Trudel et al., 2017) or 95% confidence intervals (CI) (Culliton et al. 2021) instead of SDs, the SD was derived using Cochrane-recommended formulas (Higgins et al. 2019). Where results were presented separately for males and females (Horeau et al. 2022, 2024), means and SDs were pooled using the participant-weighted method described in the Cochrane Handbook (Higgins et al. 2019).

In some studies, both groups were exposed to simulated microgravity, but one group also received an additional intervention (e.g., nutrition, artificial gravity, or thigh cuffs). Since all included studies directly reported that these interventions had no significant effect on hematological outcomes, if the data weren’t presented pooled (Nay et al. 2020), we pooled them into a single group. Means and standard deviations were combined using Cochrane-recommended formulas (Higgins et al. 2019).

Numerical values that weren’t available in text or tables were digitized from figures using WebPlotDigitizer. All hematological outcomes were converted to uniform units (e.g., hemoglobin in g/dL, serum iron in µmol/L), allowing for the use of mean differences (MD) rather than standardized mean differences (SMD), which provided more clinically interpretable effect estimates.

In the study by Dunn et al. (1984), outcome data were extracted only from the group that underwent 7 days of continuous horizontal or 6-degree head-down tilt (n = 18). The other group that completed 28 days of strict horizontal bed rest (n = 6) was excluded, as horizontal bed rest is no longer considered a valid analogue for simulated microgravity (Watenpaugh 2016). The authors reported results for the 18 participants as a single group without differentiating between horizontal and HDT positions, so we used the aggregated means and standard deviations provided for this combined group (Dunn et al. 1984).

Also, the study by Dunn et al. (1984) was excluded from the quantitative synthesis of erythropoietin (EPO) outcomes because of problems with the methods used. In that study, researchers used a fetal mouse liver cell (FMLC) bioassay to check the levels of EPO, which is known to give values that are much higher and not standardized compared to newer immunoassays like ELISA or radioimmunoassay (RIA) (Sakata et al. 1999). Since there is no reliable method to calibrate the outcomes, including this study would have resulted in less accurate pooled estimates and made comparisons between studies more difficult (Dunn et al. 1984).

In the study by Gunga et al. (1996), participants were studied under multiple conditions, including actual spaceflight and several ground-based microgravity analogues, such as −6° head-down tilt (HDT) and horizontal bed rest. Only the HDT data were used in this meta-analysis, as the authors reported that HDT was the analogue that most closely matched the response seen during real spaceflight (Gunga et al. 1996).

In the study by Trudel et al. (2017), there was no clear report of indirect (unconjugated) bilirubin. However, there were values for both total and direct (conjugated) bilirubin. So, following standard clinical practice, indirect bilirubin was calculated by subtracting direct bilirubin from total bilirubin (Trudel et al., 2017). This method is widely used to estimate unconjugated bilirubin when it is not directly given, and it is described in major clinical chemistry references such as Tietz Textbook of Clinical Chemistry (Jung 2008).

### 2.7 Risk of Bias Assessment

Risk of bias was evaluated by two reviewers using tools selected to match each study’s design and analytic structure. Because the main goal of this review was to measure changes in hematological markers after being exposed to simulated microgravity, most studies were evaluated using the NIH Quality Assessment Tool for Before–After (Pre-Post) Studies with No Control Group (National Heart Lung and Institute 2014). This choice was appropriate because, in many randomized trials, both groups underwent simulated microgravity, and the randomized co-intervention (e.g., nutritional modifications, thigh cuffs) was reported to have no measurable effect on the outcomes of interest.

One study included a non-randomized control group that did not experience simulated microgravity; it was assessed with the Joanna Briggs Institute (JBI) Checklist for Quasi-Experimental Studies (Joanna Briggs Institute 2020), which is better for comparing groups without randomization.

Each item in the selected tools was evaluated as Yes, No, Cannot Determine, Not Reported, or Not Applicable, based on the extent to which the study met the requirement.

### 2.8 Statistical Analysis

One author has performed data analysis, and it was revised by another author. It was done by Comprehensive Meta-Analysis software, version 3.0.

Pooled effect sizes were calculated as mean differences (MD) with corresponding 95% confidence intervals (CI), using the inverse-variance method for weighting. The pre-post correlation coefficient (r), required for within-subject comparisons, was neither reported in any of the included studies nor could it be calculated from the available data. Accordingly, we assumed a value of r = 0.5, as recommended by the Cochrane Handbook for Systematic Reviews of Interventions (Higgins et al. 2019). To evaluate the validity of this assumption, sensitivity analyses were conducted using alternative r values of 0.2, 0.7, and 0.9. All of these values gave us consistent estimates, with no significant changes in direction or magnitude.

Heterogeneity was evaluated using Cochran’s Q test and quantified by the I² statistic. When heterogeneity was present (p < 0.1 for Q or I² > 50%), a random-effects model was used. Without it, a fixed-effect model was used.

### 2.9 Reporting Bias Assessment

Assessment of reporting bias (e.g., by funnel plots or Egger’s test) was not performed due to the small number of studies contributing to each outcome (range: 3 to 8). The Cochrane Handbook for Systematic Reviews of Interventions says that these methods are usually not reliable and don’t have enough power when fewer than 10 studies are included in a meta-analysis (Higgins et al. 2019). For this reason, the risk of reporting bias due to missing results could not be formally assessed.

## 3. Results

### 3.1 Study Selection

The initial literature search through the previously mentioned databases has revealed 1274 articles, where 193 of them were duplicates. The number of articles identified through each database was as follows: 347 in PubMed, 19 in Cochrane, 328 in WOS, and 580 in Scopus. Title and abstract screening were done on 1081 articles, and 52 articles were retrieved, which were then screened by the full text. The authors have excluded 42 articles for the following reasons: narrative reviews (n = 5), non-human studies (n = 11), different intervention (n = 4), different outcomes (n = 7), incompatible study design (n = 5), lack of quantitative data (n = 6), insufficient duration of microgravity exposure (n = 1), mathematical modeling (n = 1), and language restrictions (n = 2).

Finally, 10 studies were eligible for inclusion in our study (Branch et al., 1998; Culliton et al., 2021; Dunn et al., 1984; Gunga et al., 1996; Horeau et al., 2022, 2024; Lampe et al., 1992; Nay et al., 2020; Ryan et al., 2016; Trudel et al., 2017). The study flow diagram is shown in **Fig. 1**.

**Fig. 1.**
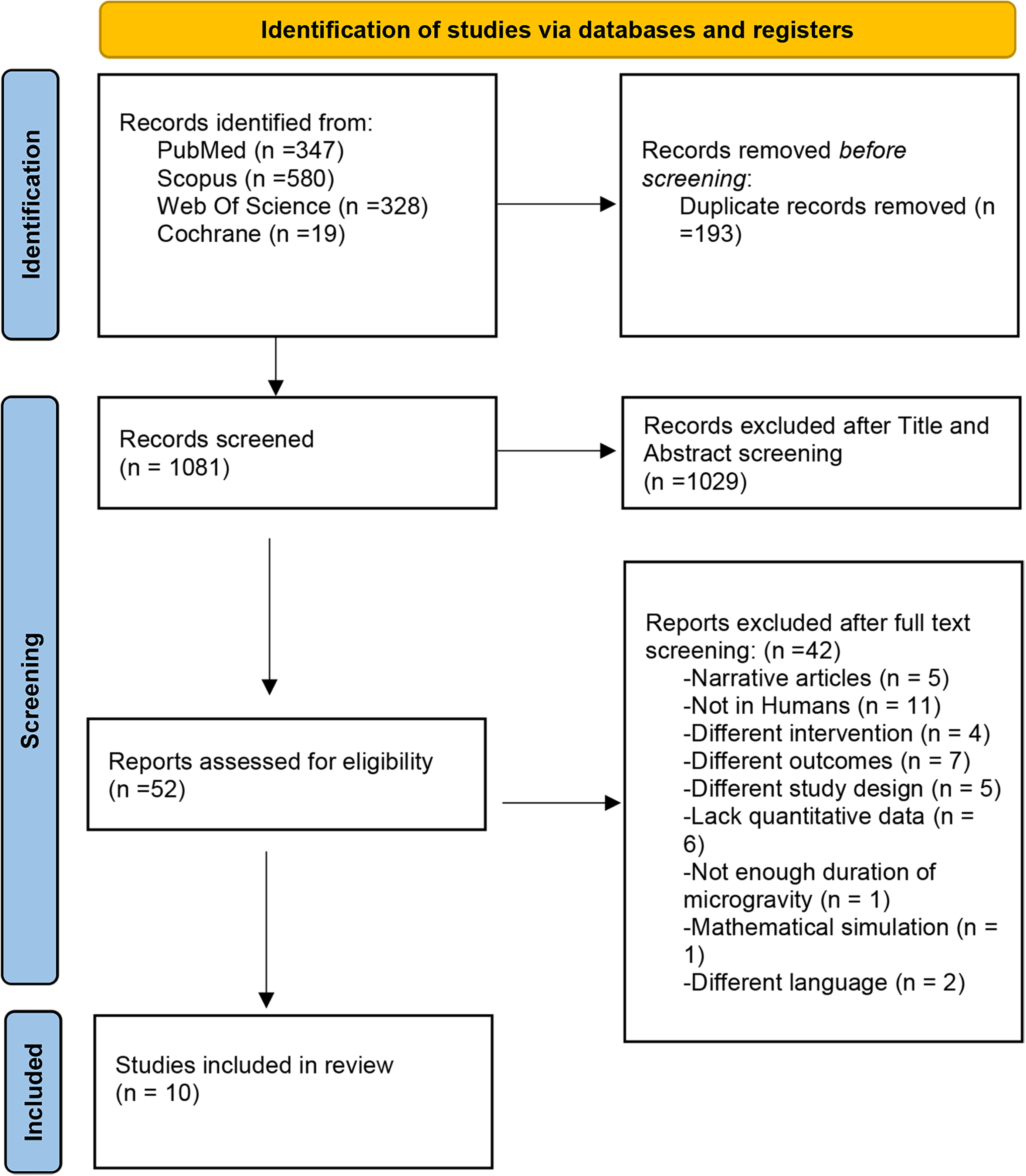
PRISMA 2020 flow diagram illustrating the study selection process.

### 3.2 Study Characteristics

Ten studies were included in this systematic review and meta-analysis, with a total of 152 participants.

Two ground-based models were used to simulate microgravity conditions. Eight studies used the 6-degree head-down tilt (HDT) model, with exposure times ranging from 4 to 60 days. Two studies used dry immersion (DI) protocols for 5 days each. Study designs included randomized clinical trials, non-randomized controlled studies, and pre-post experimental designs. Four studies included additional interventions like nutritional supplements, anti-gravity (AG) training, or thigh cuffs. The other studies did not include any additional interventions. **Table 1** displays the characteristics of the included studies. Details of the demographics of the participants are shown in **Table 2**. Reported ages ranged from the early twenties to late forties, and body mass index (BMI) values were generally within normal limits. Most participants were male, although two studies included female participants. In general, the studies were very heterogeneous in terms of methodology (design, type of intervention, and characteristics of participants), which provided a broad dataset to assess the physiological impact of simulated microgravity. Baseline values for the measured physiological outcomes are presented in **Table 3**.

**Table 1.** Characteristics of included studies.

| Study Name | Study Design | Microgravity Type | Exposure Duration | Sample Size (n) | Participant Gender | Microgravity Control Group Present? | Other Intervention Control Group Present? | Intervention Type |
| --- | --- | --- | --- | --- | --- | --- | --- | --- |
| Branch1998 | Non-randomized Controlled Experimental Study | 6-degree head-down tilt (HDT) | 16 days | 6 | 6 males | Yes | No |  |
| Culliton2021 | Randomized Clinical Trial | 6-degree head-down tilt (HDT) | 60 days | 20 | 20 males | No | Yes | Nutrition |
| dunn1984 | Experimental Study - Pre-post Design | Continuous horizontal or 6-degree head-down tilt (HDT) | 7 days | 18 | 18 males | No | No |  |
| gunga1996 | Experimental Study - Pre-post Design | 6-degree head-down tilt (HDT) | 42 days | 8 | 8 males | No | No |  |
| Horeau2022 | Randomized Clinical Trial | 6-degree head-down tilt (HDT) | 60 days | 23 | 7 females, 16 males | No | Yes | AG training |
| Horeau2024 | Experimental Study - Pre-post Design | Dry Immersion (DI) | 5 days | 37 | 18 females, 19 males | No | No |  |
| lampe1992 | Experimental Study - Pre-post Design | 6-degree head-down tilt (HDT) | 10 days | 6 | 6 males | No | No |  |
| Nay 2020 | Randomized Clinical Trial | Dry Immersion (DI) | 5 days | 17 | 18 males | No | Yes | Thigh cuffs |
| ryan2016 | Experimental Study - Pre-post Design | 6-degree head-down tilt (HDT) | 4 days | 7 | 7 males | No | No |  |
| trudel2017 | Cross-over Experimental Study | 6-degree head-down tilt (HDT) | 21 days | 10 | 10 males | No | Yes | Nutrition |

\*AG: Artificial Gravity

**Table 2.** Baseline demographic characteristics of participants in the included studies.

| Study Name | Age (years) | BMI (kg/m<sup>2</sup>) | Height (cm) | Weight (kg) | VO<sub>2</sub> Max (ml/min/kg) |
| --- | --- | --- | --- | --- | --- |
| Branch1998 | 39 ± 7 |  | 182.0 ± 6.5 | 81.7 ± 12.0 | 32.7 ± 8.1 |
| Culliton2021 | 34.2 ± 7.8 | 22–27 |  |  |  |
| dunn1984 |  |  |  | 68.8 ± 2.12 |  |
| gunga1996 | 27.9 ± 2.3 |  | 175 ± 4 | 74.5 ± 8.3 |  |
| Horeau2022 | 33.27 ± 9.30 | 24.30 ± 1.99 | 174.03 ± 5.91 | 74.57 ± 7.83 |  |
| Horeau2024 | 28.38 ± 4.51 | 22.46 ± 1.83 | 170.83 ± 5.06 | 65.82 ± 6.49 |  |
| lampe1992 | 26 |  |  |  | 42.5 |
| Nay 2020 | 33.6 ± 5.5 | 23.5 ± 2.5 | 178 ± 6 | 74.4 ± 8.0 | 46.1 ± 6.9 |
| ryan2016 | 21 ± 3 |  | 180 ± 11 | 73 ± 10 | 50 ± 6 |
| trudel2017 | 32 | 23.2 |  |  |  |

Age, body mass index (BMI), height, weight, and VO<sub>2</sub> max are reported as mean ± standard deviation.

**Table 3.** Baseline values for the measured physiological outcomes across included studies.

| Study Name | Hemoglobin (g/dL) | Hematocrit (%) | Erythropoietin (mIU/ml) | Plasma Volume (ml) | RCV (ml) | RBC Count (×10<sup>6</sup>/μL) | Reticulocytes (%) | Total Hemoglobin Mass (g) | MCV (fL) | MCH (pg) | MCHC (g/dL) | Iron (μmol/L) | Transferrin Saturation (%) | Bilirubin (μmol/L) | Indirect Bilirubin (μmol/L) | Haptoglobin (g/L) |
| --- | --- | --- | --- | --- | --- | --- | --- | --- | --- | --- | --- | --- | --- | --- | --- | --- |
| Branch1998 | 14.5 ± 0.9 | 43.0 ± 3.4 | 11.6 ± 2.9 | 3769 ± 512 | 2845 ± 410 |  |  |  |  |  |  |  |  |  |  |  |
| Culliton2021 | 13.7 ± 0.64 |  | 12.2 ± 4.3 |  |  | 4.529 ± 0.29 | 1.05 ± 0.39 | 818.2 ± 104.6 |  |  |  | 14.57 ± 3.47 | 29.0 ± 9.1 | 11.5 ± 4.8 |  | 1.03 ± 0.29 |
| dunn1984 |  | 41.4 ± 2.12 | 120 ± 84.9 |  |  |  | 0.60 ± 0.3 |  |  |  |  |  |  |  |  |  |
| gunga1996 | 14.2 ± 1.4 | 44.2 ± 2.2 | 11.4 ± 3.3 |  |  |  |  |  |  |  |  |  |  |  |  |  |
| Horeau2022 | 14.96 ± 1.01 | 39.75 ± 4.82 |  | 3030.61 ± 480.49 | 2382.35 ± 374.77 | 4.69 ± 0.35 | 0.95 ± 0.19 | 813.59 ± 122.45 | 86.39 ± 4.38 | 29.7 ± 1.66 | 34.39 ± 0.64 | 16.35 ± 4.05 | 26.33 ± 6.45 | 11.74 ± 6.53 | 11.40 ± 6.61 | 1.01 ± 0.40 |
| Horeau2024 |  |  |  | 3275.76 ± 571.05 |  | 4.64 ± 0.29 | 1.02 ± 0.35 | 710.46 ± 103.14 | 91.49 ± 3.52 | 31.11 ± 1.1 | 34.11 ± 0.40 | 18.08 ± 1.82 | 30.49 ± 2.96 |  |  |  |
| lampe1992 | 14.5 ± 1.10 | 42.3 ± 3.46 |  |  |  | 4.73 ± 0.39 |  |  | 90.5 ± 2.06 | 31.4 ± 1.57 | 34.6 ± 1.07 |  |  |  |  |  |
| Nay 2020 | 14.55 ± 0.88 | 41.96 ± 2.33 | 7.70 ± 2.70 |  |  | 4.85 ± 0.37 | 1.15 ± 0.20 |  |  |  |  | 19.06 ± 2.74 | 32.69 ± 4.90 | 8.00 ± 2.27 | 4.65 ± 1.57 | 0.85 ± 0.23 |
| ryan2016 | 15.7 ± 0.4 | 45.3 ± 1.8 | 7.1 ± 1.8 | 3369 ± 569 | 2356 ± 371 |  | 1.33 ± 0.28 | 817 ± 135 |  |  | 34.7 ± 1.0 |  |  | 8.55 ± 4.1 |  | 0.54 ± 0.32 |
| trudel2017 | 14.0 ± 0.32 | 41.1 ± 1.3 | 10.8 ± 2.85 | 4004 ± 351 | 2459 ± 196.1 | 4.83 ± 0.16 | 1.10 ± 0.14 | 838 ± 66.4 | 83.9 ± 1.3 | 28.9 ± 0.3 | 34.1 ± 0.3 |  |  | 14.5 ± 4.3 | 8.9 ± 4.55 | 1.11 ± 0.44 |

Values are presented as mean ± standard deviation

### 3.3 Risk of Bias Assessment

Risk of bias was assessed using two tools based on study design. Nine studies were assessed by the NIH Quality Assessment Tool for Before-After (Pre-Post) Studies without Control Groups, and one study was assessed by the JBI Checklist for Quasi-Experimental Studies. Low to moderate risk of bias was found in the majority of the included studies. For the studies that were evaluated by the NIH Tool, the majority met the criteria for clearly defined research questions, valid outcome measurements, and appropriate statistical analyses. However, due to the specialized nature of this field of study, sample sizes were generally small, and several studies lacked clear details regarding blinding procedures. **Table 4** provides a detailed overview of the risk of bias evaluations for every study.

**Table 4.** Risk of bias assessment using NIH Tool for Before-After (Pre-Post) Studies with No Control Group.

| Study | Q1 | Q2 | Q3 | Q4 | Q5 | Q6 | Q7 | Q8 | Q9 | Q10 | Q11 | Q12 |
| --- | --- | --- | --- | --- | --- | --- | --- | --- | --- | --- | --- | --- |
| **Culliton et al., 2021** | Y | Y | Y | NR | Y | Y | Y | NR | Y | Y | Y | NA |
| **Dunn et al., 1984** | Y | N | CD | NR | Y | Y | Y | NR | Y | Y | Y | NA |
| **Gunga et al., 1996** | Y | Y | Y | NR | CD | Y | Y | NR | Y | Y | Y | NA |
| **Horeau et al., 2022** | Y | Y | Y | NR | Y | Y | Y | NR | Y | Y | Y | NA |
| **Horeau et al., 2024** | Y | Y | Y | NR | Y | Y | Y | NR | Y | Y | N | NA |
| **Lampe et al., 1992** | Y | Y | CD | NR | CD | Y | Y | NR | Y | Y | Y | NA |
| **Nay et al., 2020** | Y | Y | Y | NR | Y | Y | Y | NR | Y | Y | N | NA |
| **Ryan et al., 2016** | Y | Y | Y | NR | CD | Y | Y | NR | Y | Y | Y | NA |
| **Trudel et al., 2017** | Y | Y | Y | NR | CD | Y | Y | NR | Y | Y | Y | NA |

\*Y, yes; N, No; CD, cannot determine; NA, not applicable; NR, not reported

For the study that was evaluated by the JBI Tool, most criteria were met, including clarity of cause-effect relationships, similarity between groups, and appropriate statistical methods, but the study did not include multiple pre- and post-intervention measurements except for the erythropoietin, which might be considered a drawback. Details of the assessment are shown in **Table 5**.

**Table 5.** Risk of bias assessment using JBI Checklist for Quasi-Experiments for the study Branch et al., 1998.

| **JBI Checklist Item** | **Assessment** |
| --- | --- |
| **1. Is it clear what is the 'cause' and what is the 'effect'?** | Yes |
| **2. Were the participants in any comparisons similar?** | Yes |
| **3. Were the participants receiving similar care other than the exposure?** | Yes |
| **4. Was there a control group?** | Yes |
| **5. Were there multiple measurements of the outcome pre/post intervention?** | No |
| **6. Was follow-up complete and adequately described?** | Yes |
| **7. Were outcomes measured the same way in both groups?** | Yes |
| **8. Were outcomes measured in a reliable way?** | Yes |
| **9. Was appropriate statistical analysis used?** | Yes |

### 3.4 Outcomes

#### 3.4.1 Core Anemia and Circulatory Volume Indicators Hemoglobin

Eight studies were analyzed to assess the changes in hemoglobin (Hb) levels during the exposure to microgravity. The meta-analysis showed a statistically significant increase in Hb concentrations with a consistent elevation across studies, which was indicated by mean difference (MD). The pooled value was +1.21 g/dL (95% CI: 0.86 to 1.57, p < 0.001). The random-effects model was used because of the significant heterogeneity (I² = 71.85%, p = 0.001) **(Fig. 2a)**.

**Fig. 2.**
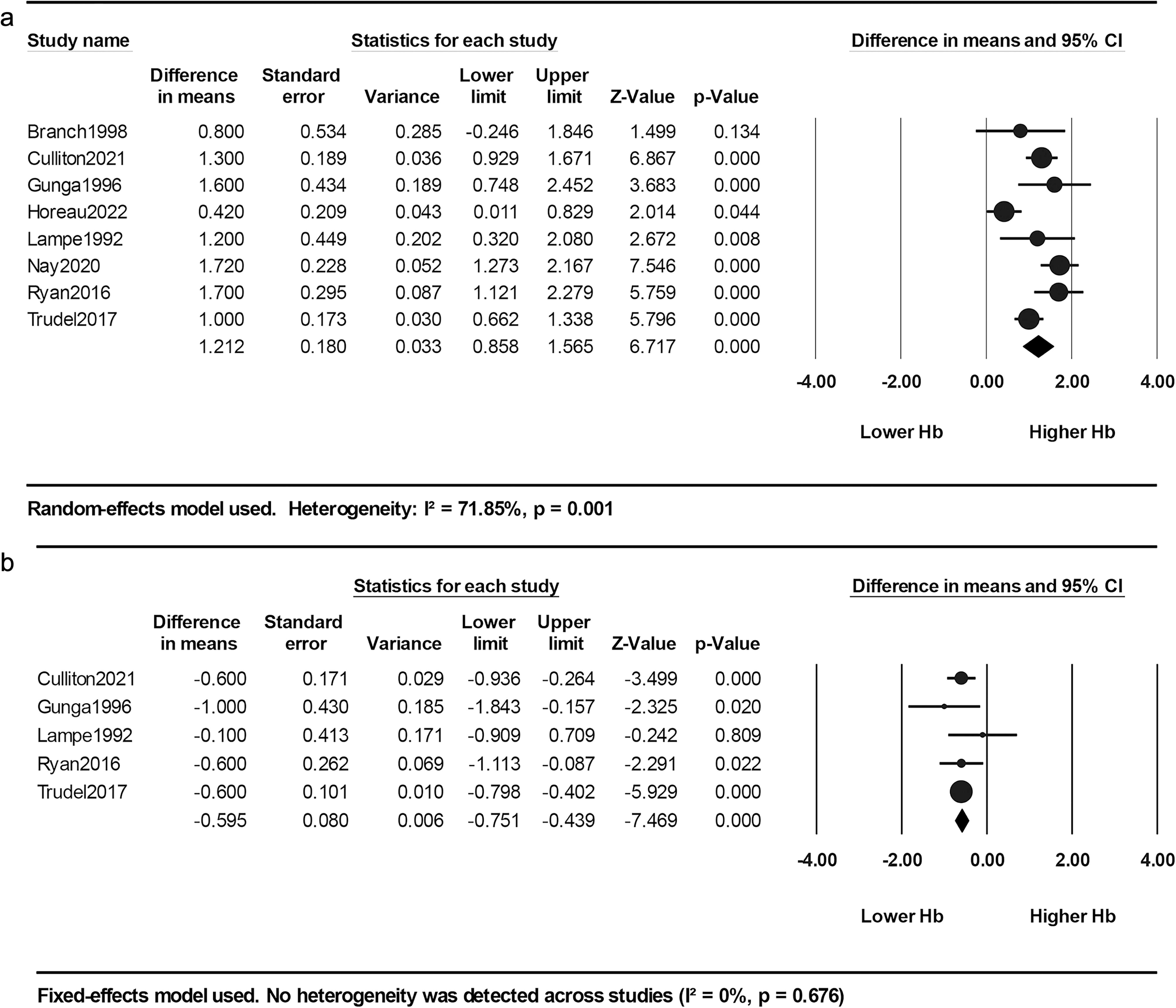
Forest plots of changes in Hemoglobin concentrations (a) Hemoglobin during exposure to simulated microgravity (b) Hemoglobin after reambulation.

Five studies were analyzed to assess the changes after reambulation to normal gravity. In contrast, the meta-analysis showed a significant reduction in Hb levels. The pooled MD value was –0.60 g/dL (95% CI: –0.75 to –0.44, p < 0.001). As no heterogeneity was detected (I² = 0%, p = 0.676), the fixed-effects model was used **(Fig. 2b)**.

##### Hematocrit

Eight studies were analyzed to assess the changes in hematocrit (Hct) levels during the exposure to microgravity. The meta-analysis showed a statistically significant increase in Hct concentrations, which was reflected by the mean difference (MD). The pooled value was +3.10% (95% CI: 1.85 to 4.34, p < 0.001). The random-effects model was used because of the significant heterogeneity (I² = 82.27%, p < 0.001) **(Fig. 3a)**.

**Fig. 3.**
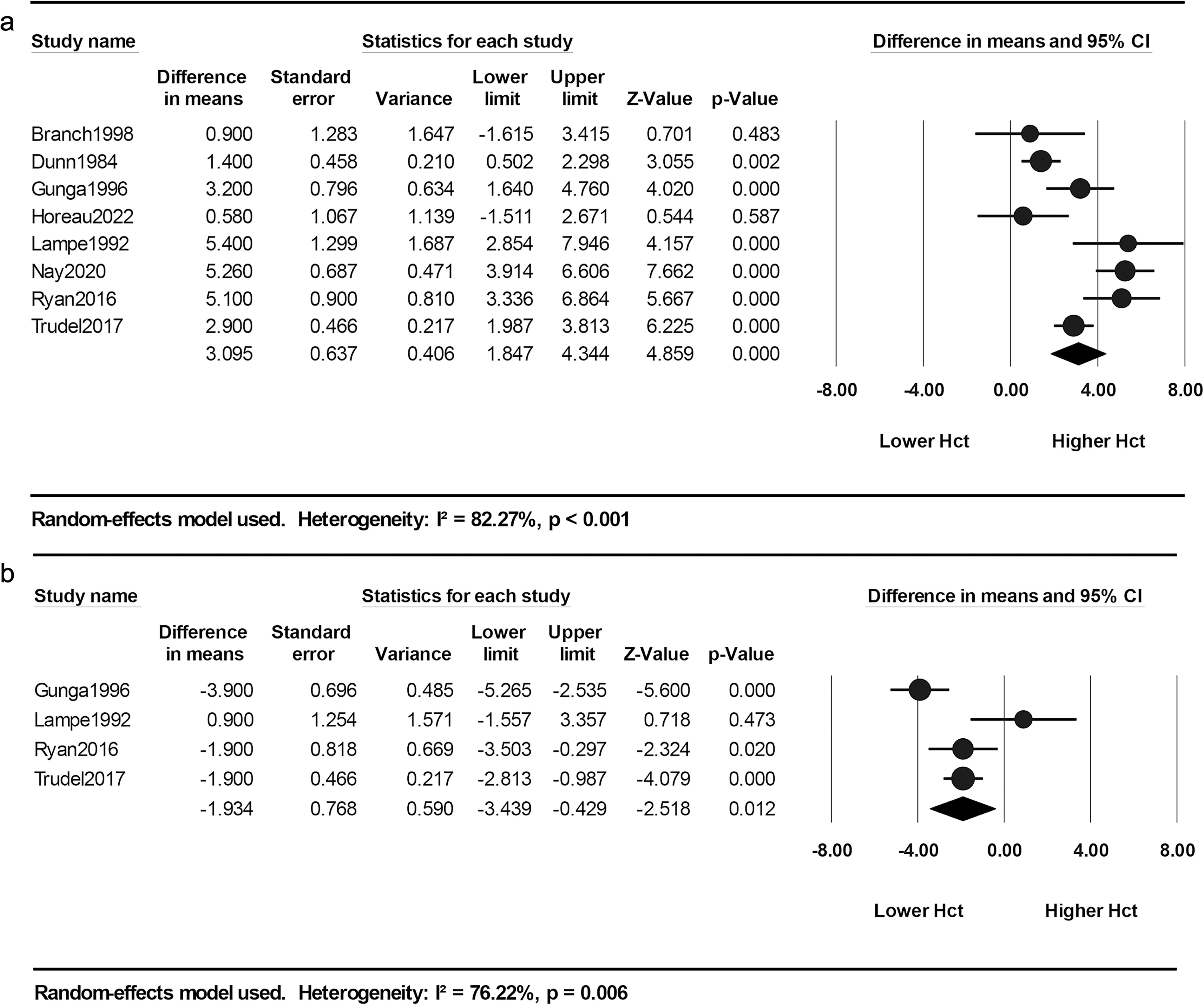
Forest plots of changes in Hematocrit concentrations (a) Hematocrit during exposure to simulated microgravity (b) Hematocrit after reambulation.

After reambulation to normal gravity, Four studies were analyzed to assess the changes in Hct levels. The meta-analysis showed a significant reduction in Hct levels. The pooled MD value was –1.93% (95% CI: –3.44 to –0.43, p = 0.012). Because of the significant heterogeneity (I² = 76.22%, p = 0.006), a random-effects model was used **(Fig. 3b)**.

##### Total Hemoglobin Mass

Five studies were used to assess the changes in total hemoglobin mass (tHb mass) during the exposure to microgravity. The meta-analysis showed a statistically significant reduction in tHb mass, which was reflected by mean difference (MD). The pooled value was –58.06 g (95% CI: –88.94 to –27.17, p < 0.001). The random-effects model was used because of the moderate heterogeneity (I² = 51.81%, p = 0.081) **(Fig. 4a)**.

**Fig. 4.**
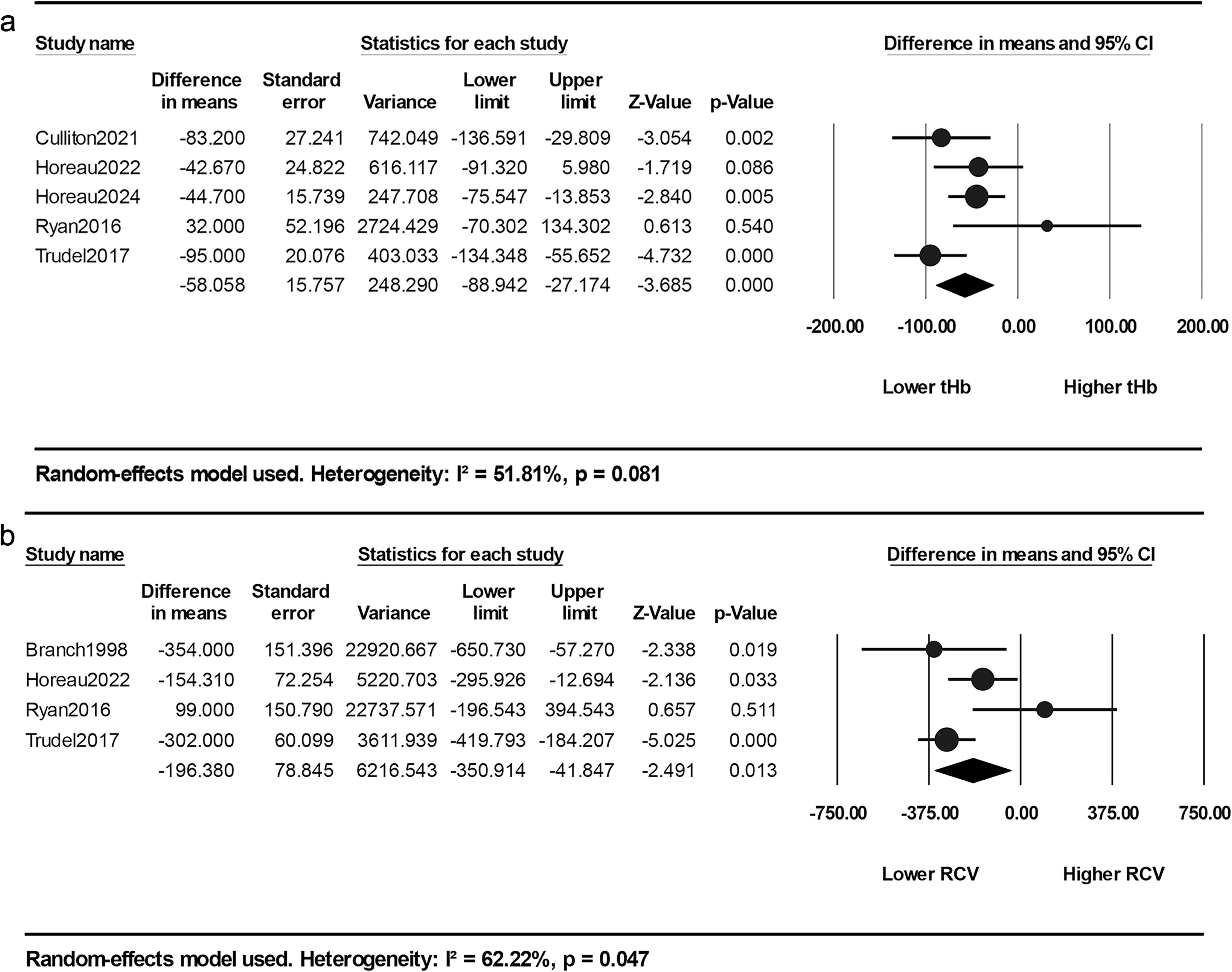
Forest plot of changes in (a) total hemoglobin mass and (b) red cell volume during simulated microgravity exposure.

##### Red Cell Volume

Four studies were analyzed to assess the changes in red cell volume (RCV) during the exposure to microgravity. The meta-analysis showed a statistically significant decrease in RCV, which was demonstrated by mean difference (MD). The pooled value was –196.38 mL (95% CI: –350.91 to –41.85, p = 0.013). The random-effects model was used because of the moderate heterogeneity (I² = 62.22%, p = 0.047) **(Fig. 4b)**.

##### Plasma Volume

Five studies were used to assess the changes in plasma volume (PV) during the exposure to microgravity. The analysis showed a statistically significant reduction in PV, which was reflected by mean difference (MD). The pooled value was –533.10 mL (95% CI: –722.08 to –344.13, p < 0.001). The random-effects model was used because of the high heterogeneity (I² = 69.87%, p = 0.010) **(Fig. 5)**.

**Fig. 5.**
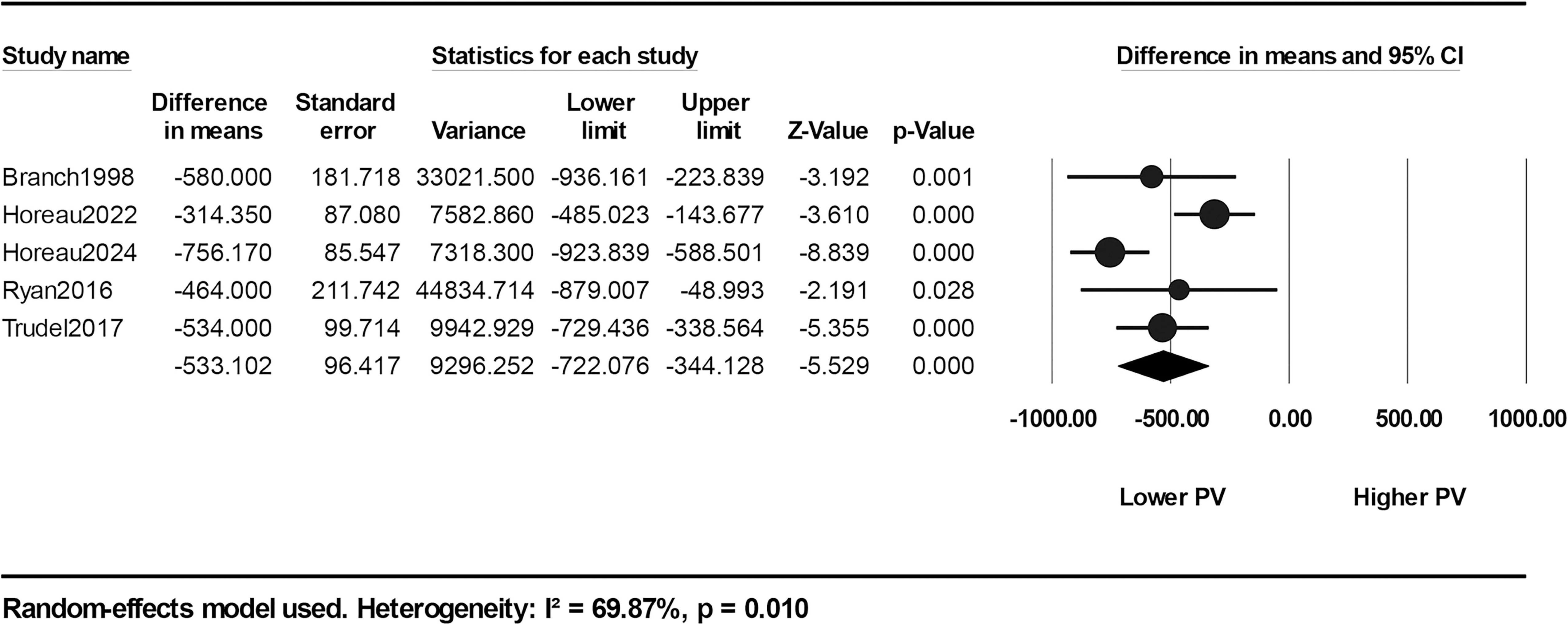
Forest plot of changes in plasma volume during simulated microgravity exposure.

#### 3.4.2 Erythropoiesis and Red Cell Indices Erythropoietin

Six studies were analyzed to assess the changes in erythropoietin (EPO) levels during the exposure to microgravity. The meta-analysis showed a statistically significant decline in EPO concentrations, which was demonstrated by mean difference (MD).

The pooled value was –1.96 mIU/mL (95% CI: –2.65 to –1.26, p < 0.001). As low heterogeneity was detected (I² = 22.3%, p = 0.266), the fixed-effects model was used. **(Fig. 6a)**.

**Fig. 6.**
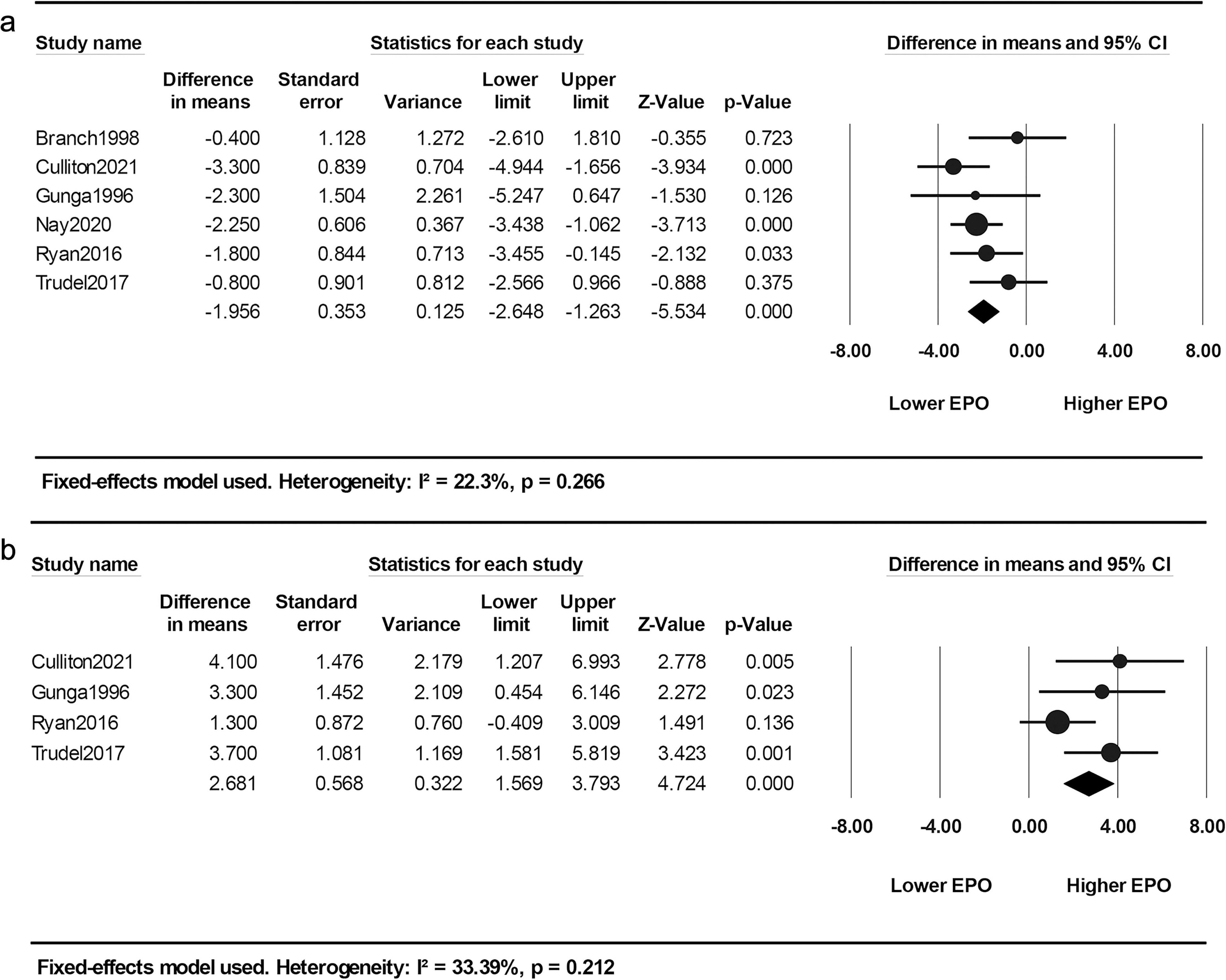
Forest plots of changes in erythropoietin concentration (a) during simulated microgravity exposure and (b) after reambulation.

After reambulation to normal gravity, Four studies were analyzed to assess the changes in EPO levels. The meta-analysis showed a significant rebound in EPO levels. The pooled MD value was +2.68 mIU/mL (95% CI: 1.57 to 3.79, p < 0.001). The fixed-effects model was used as there was low heterogeneity (I² = 33.39%, p = 0.212) **(Fig. 6b)**.

##### Reticulocyte Count

Six studies were used to assess the changes in reticulocyte percentage during the exposure to microgravity. The meta-analysis indicated no statistically significant change, which was demonstrated by mean difference (MD). The pooled value was – 0.05% (95% CI: –0.15 to 0.04, p = 0.279). The random-effects model was used because of the significant heterogeneity (I² = 71.37%, p = 0.004) **(Fig. 7a)**.

**Fig. 7.**
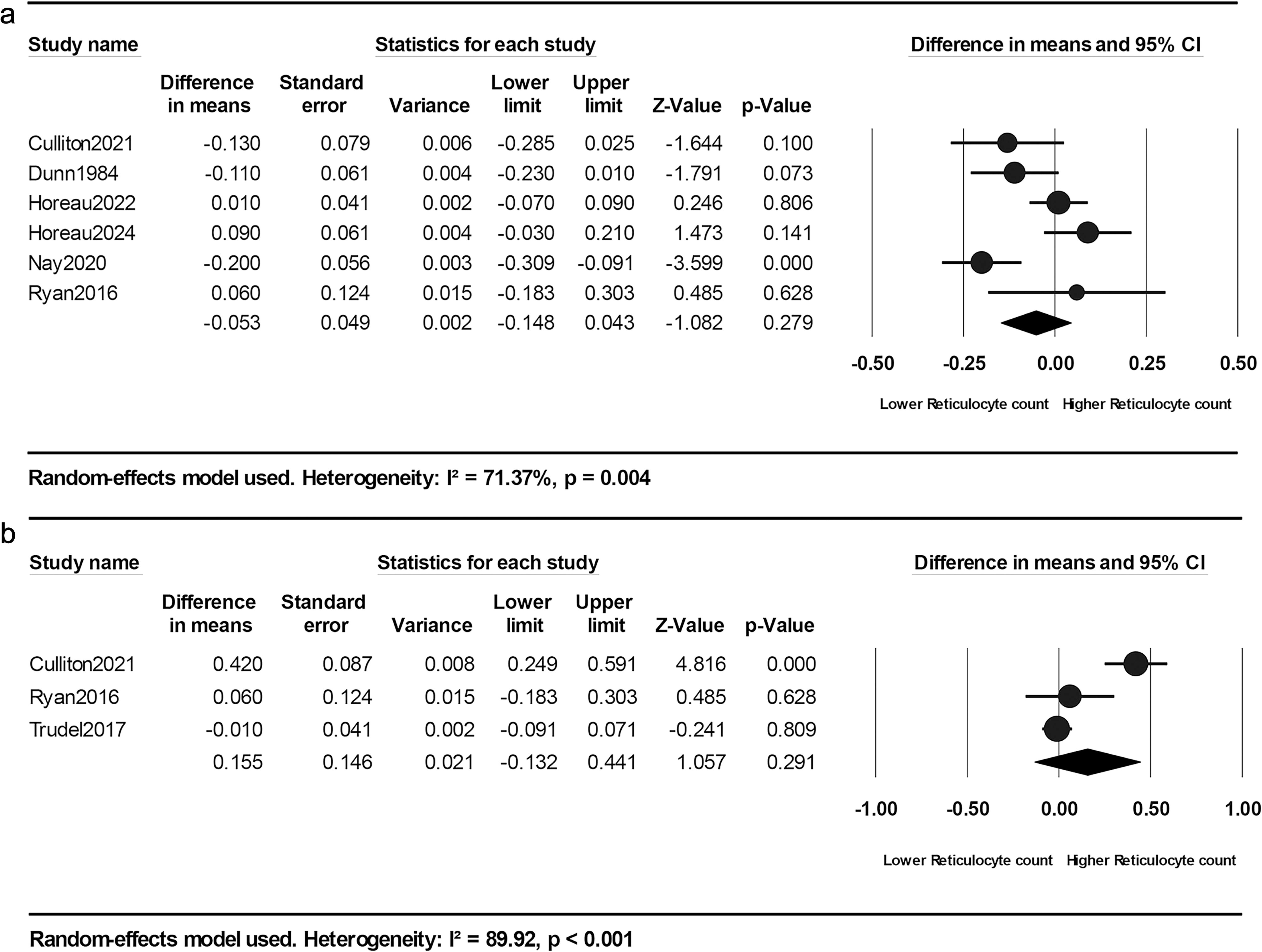
Forest plots of changes in reticulocyte percentage (a) during simulated microgravity exposure and (b) after reambulation.

After returning to normal gravity, three studies were analyzed to assess the changes in reticulocyte percentage. Although a slight increase was observed (MD = +0.16%, 95% CI: –0.13 to 0.44), this difference was not statistically significant (p = 0.291). A random-effects model was used as there was high heterogeneity (I² = 89.92%, p < 0.001) **(Fig. 7b)**.

##### Red Blood Cell Count

Six studies were analyzed to assess the changes in red blood cell (RBC) count during the exposure to microgravity. The meta-analysis showed a statistically significant increase in RBC levels, which was reflected by the mean difference (MD). The pooled value was +0.41 × 10⁶/µL (95% CI: 0.24 to 0.58, p < 0.001). The random-effects model was used because of the significant heterogeneity (I² = 88.16%, p < 0.001) **(Fig. 8a)**.

**Fig. 8.**
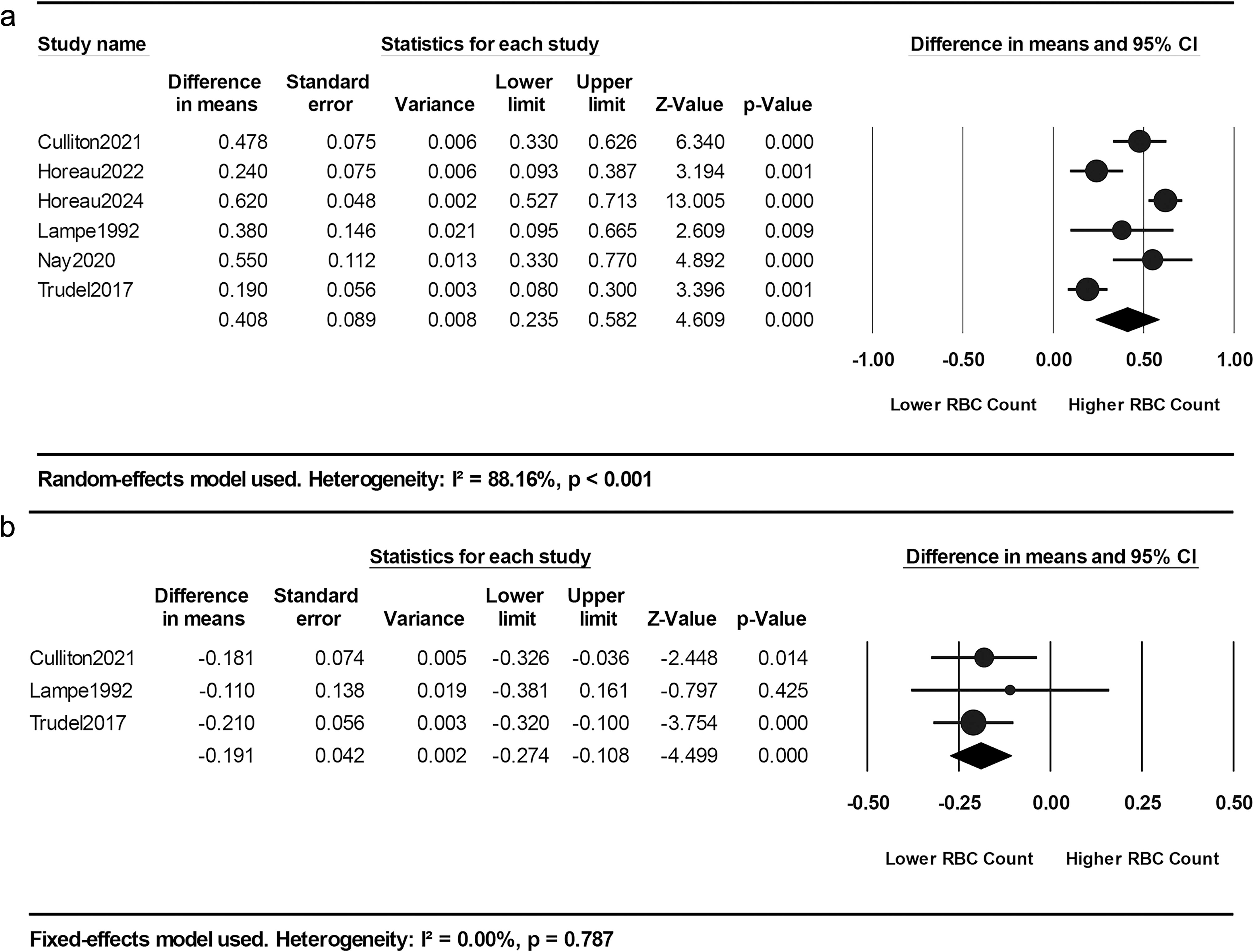
Forest plots of changes in red blood cell count (a) during simulated microgravity exposure and (b) after reambulation.

After returning to normal gravity, three studies were analyzed to assess the changes in RBC levels. The meta-analysis showed a significant decrease in RBC levels. The pooled MD value was –0.19 × 10⁶/µL (95% CI: –0.27 to –0.11, p < 0.001). No heterogeneity was detected (I² = 0%, p = 0.787), so a fixed-effects model was applied **(Fig. 8b)**.

#### 3.4.3 Red Cell Morphological Indices Mean Corpuscular Volume

Four studies were analyzed to assess the changes in mean corpuscular volume (MCV) during the exposure to microgravity. The meta-analysis indicated no statistically significant change, which was demonstrated by mean difference (MD). The pooled value was +0.13 fL (95% CI: –0.48 to 0.74, p = 0.673). The fixed-effects model was used because of the moderate heterogeneity (I² = 46.24%, p = 0.134) **(Supplementary Fig. 1)**.

##### Mean Corpuscular Hemoglobin

Four studies were used to assess the changes in mean corpuscular hemoglobin (MCH) during the exposure to microgravity. The meta-analysis indicated no statistically significant change, which was evidenced by mean difference (MD). The pooled value was +0.08 pg (95% CI: –0.14 to 0.30, p = 0.482). A fixed-effects model was used because of the absence of heterogeneity (I² = 0%, p = 0.970) **(Supplementary Fig. 2)**.

##### Mean Corpuscular Hemoglobin Concentration

Five studies were used to assess the changes in mean corpuscular hemoglobin concentration (MCHC) during the exposure to microgravity. The pooled mean difference was +0.25 g/dL (95% CI: –0.03 to 0.53, p = 0.084), which reflects no statistically significant change. The random-effects model was used because of the high heterogeneity (I² = 77.37%, p = 0.001) **(Supplementary Fig. 3)**.

#### 3.4.4 Iron Metabolism and Hemolysis-Associated Biomarkers Serum Iron

Four studies were analyzed to assess the changes in serum iron concentrations during the exposure to microgravity. The meta-analysis showed a statistically significant increase in iron levels, which was reflected by the mean difference (MD). The pooled value was +4.59 µmol/L (95% CI: 0.97 to 8.20, p = 0.013). The random-effects model was used because of the significant heterogeneity (I² = 96.2%, p < 0.001) **(Fig. 9a)**.

**Fig. 9.**
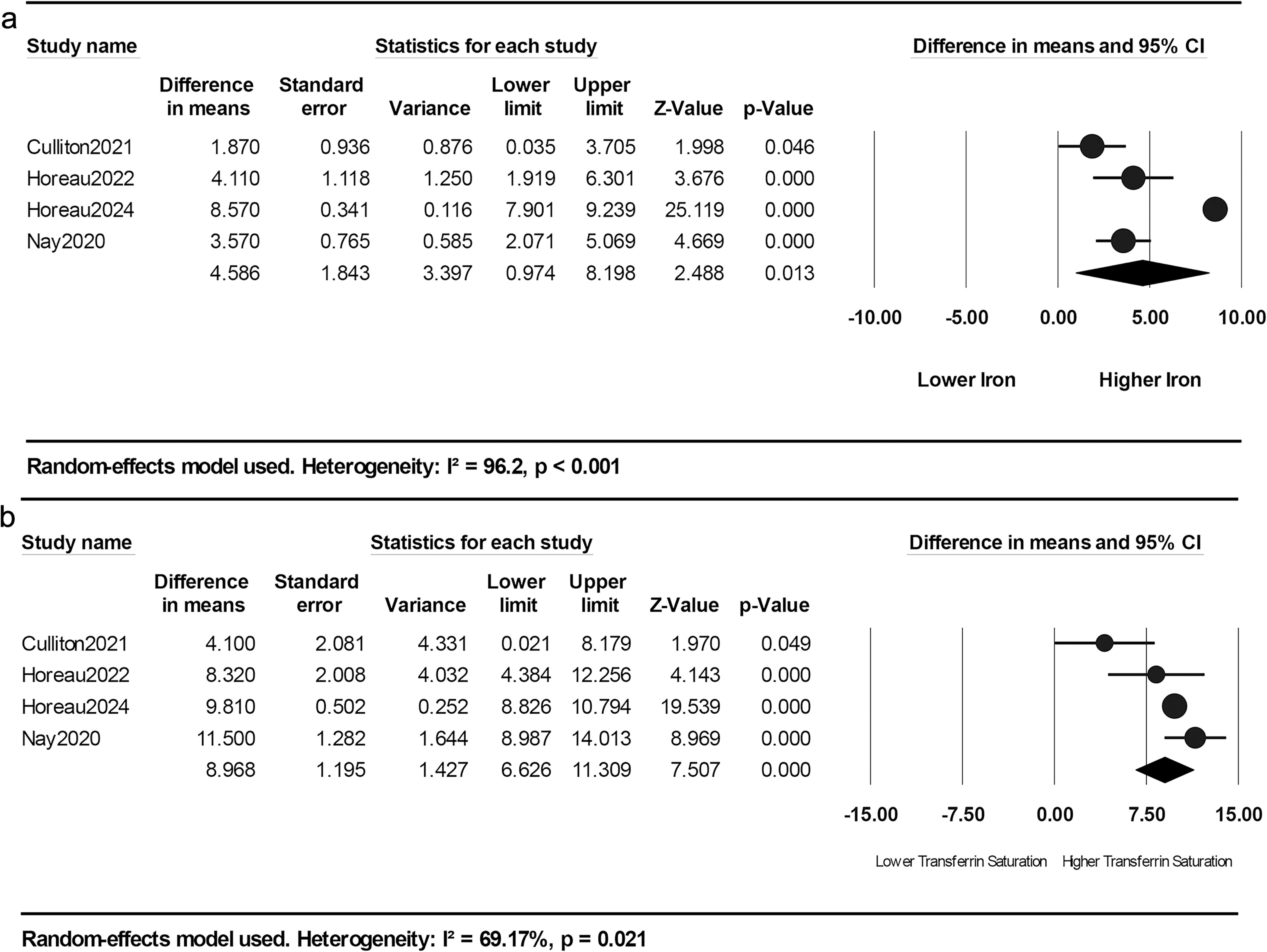
Forest plots of changes in (a) serum iron and (b) transferrin saturation during simulated microgravity exposure.

##### Transferrin Saturation

Four studies were analyzed to assess the changes in transferrin saturation during the exposure to microgravity. Meta-analysis showed a statistically significant increase, which was reflected by mean difference (MD). The pooled value was +8.97% (95% CI: 6.63 to 11.31, p < 0.001). The random-effects model was used because of the moderate heterogeneity (I² = 69.17%, p = 0.021) **(Fig. 9b)**.

##### Total Bilirubin

Five studies were used to assess the changes in total bilirubin levels during the exposure to microgravity. The meta-analysis indicated no statistically significant change, which was demonstrated by mean difference (MD). The pooled value was +0.10 µmol/L (95% CI: –1.36 to 1.56, p = 0.890). The random-effects model was used because of the moderate heterogeneity (I² = 56.78%, p = 0.055) **(Fig. 10a)**.

**Fig. 10.**
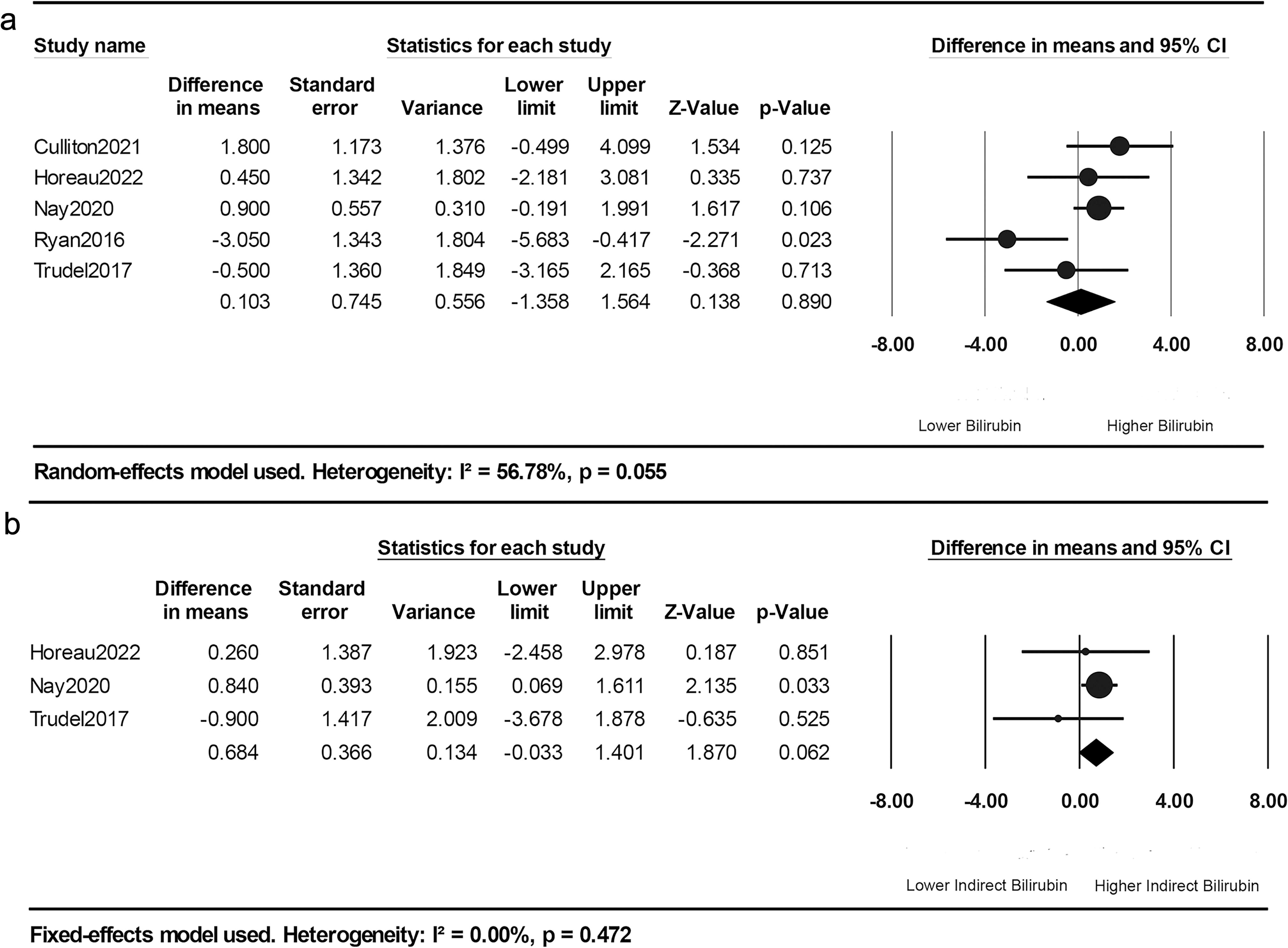
Forest plots of changes in (a) total bilirubin concentration, (b) indirect bilirubin concentration during simulated microgravity exposure.

##### Indirect Bilirubin

Three studies were used to assess the changes in indirect (unconjugated) bilirubin levels during the exposure to microgravity. The meta-analysis demonstrated no statistically significant change, which was demonstrated by mean difference (MD). The pooled value was +0.68 µmol/L (95% CI: –0.03 to 1.40, p = 0.062), which did not reach statistical significance. No heterogeneity was detected (I² = 0%, p = 0.472), so a fixed-effects model was applied **(Fig. 10b)**.

##### Haptoglobin

Five studies were used to assess the changes in haptoglobin concentrations during the exposure to microgravity. The meta-analysis indicated no statistically significant change, which was demonstrated by mean difference (MD). The pooled value was +0.063 g/L (95% CI: –0.007 to 0.133, p = 0.077). No heterogeneity was detected (I² = 0%, p = 0.500), so a fixed-effects model was applied **(Supplementary Fig. 4)**.

## 4. Discussion

This systematic review and meta-analysis dealt with data from ten studies to try to give a comprehensive overview of the hematological effects of exposure to short-to medium-duration simulated microgravity in humans. Because of the relevance of models like head-down tilt and dry immersion to both spaceflight and prolonged physical inactivity on earth, studying the effect of these models can help us understand the physiological changes that happen and then help us develop countermeasures to overcome them, whether in space or in clinical strategies for patients on earth (Hargens et al. 2013).

Analysis of various physiological biomarkers showed several effects upon exposure to microgravity, such as hemoconcentration due to plasma volume contraction, decreased erythropoiesis, iron metabolism changes, and some degree of quantitatively unsupported hemolysis based on available biomarker data. These changes appeared to occur without significant changes to the red blood cell morphology, which supports the idea that those are physiological adaptations rather than a pathological response (Smith 2002).

The early redistribution of body fluids and the resulting decrease in plasma volume (Iwase et al. 2020) are the most important physiological changes that explain many of the observed results. This causes hemoconcentration, which makes the blood look more concentrated not because an actual increase in red blood cells happened but because of the decrease in the plasma volume (Otto et al. 2017). This particular change occurs quickly after exposure to simulated microgravity, which greatly affects how several hematological markers should be interpreted.

That is why the increase in hemoglobin and hematocrit should not be seen as a true elevation of red cell mass but as a result of the plasma volume reduction (Garrett-Bakelman et al. 2019). The first observation that supports this interpretation is that after moving back to normal gravity, both hemoglobin and hematocrit levels drop significantly below baseline levels. The second observation is that total hemoglobin mass (tHb) and red cell volume (RCV), which are independent of blood volume so they are better indicator in this situation (Minetti et al. 2022), both reflect real physiological loss and decline upon exposure.

The same pattern is seen with red blood cell count, which initially seems to be elevated during exposure because of its dependence on plasma volume for concentration. However, after reambulation, this increased RBC count is reversibly decreased beyond baseline levels. This further supports that early changes are hemoconcentration, not enhanced erythropoiesis.

The actual decrease in red blood cell volume and associated indices can be linked to the fact that erythropoietic activity was low during simulated microgravity exposure. This is best reflected by the constant reduction of erythropoietin (EPO), which is an important hormone regulating red blood cell production (Jelkmann 2011). The reduction in EPO levels is probably central in mediating the decrease in total hemoglobin mass and red cell volume because erythropoietic drive is actively reduced (Keohane et al. 2020).

Notably, the EPO suppression seems to be a secondary effect of reduced plasma volume. The central fluid shift caused by microgravity raises renal perfusion and oxygenation, which are well-known causes of EPO suppression in microgravity analogs. This further shows the importance of fluid dynamics in determining how the body responds to microgravity (Garrett-Bakelman et al. 2019). It also supports the idea that these changes are a coordinated physiological adaptation rather than a pathological process.

After reambulation, EPO levels rise again, likely because the plasma volume has been restored. This hormonal recovery allows other blood parameters to gradually return to normal over time. This supports the idea that spaceflight-associated anemia is predominantly a self-resolving condition recovering gradually after landing, assuming no permanent organ damage has developed (Trudel et al., 2022).

Because of erythropoietin suppression and general downregulation of erythropoiesis, there are no significant changes in reticulocyte percentage during or post exposure to simulated microgravity. This is consistent with the assumption that there is a hypoproliferative state where there is diminished bone marrow output (Blaber et al. 2015). Moreover, the short follow-up period after reambulation in most cases probably limited the ability to observe any delayed reticulocyte rebound that may occur following the EPO recovery post-reambulation.

This suppressed erythropoietic response also makes interpreting potential hemolysis more difficult. Physiologically, reticulocyte production usually increases in response to hemolysis (van Galen and Simsek 2022; Rai et al. 2025), but in our condition that response may be muted or masked due to the existing hypoproliferative state. Therefore, not observing reticulocytosis cannot be taken as conclusive proof that there is no hemolysis, which represents a major limitation concerning the interpretation of hematological dynamics under suppressed erythropoiesis.

In all studies evaluating mean corpuscular volume (MCV), mean corpuscular hemoglobin (MCH), and mean corpuscular hemoglobin concentration (MCHC), no significant changes were noted during exposure to simulated microgravity. These indices reflect the dimensions and hemoglobin content of red blood cells (Walker et al. 1990) and, given their constancy, suggest that erythropoiesis, although decreased in quantity, still maintained some structural order, as there were no macrocytic nor microcytic changes observed.

Simulated microgravity exposure was associated with an increase in serum iron concentration, alongside a rise in transferrin saturation. Collectively, these results indicate that there is a measurable elevation in the circulating iron levels (Zwart et al. 2013). Increased circulating iron levels may result from microgravity’s hypoproliferative effects slowing the utilization of iron for erythropoiesis. It may also be due to the release of iron from red blood cells as a consequence of hemolysis. Likely, both processes, with different contributions, led to the changes observed.

In assessing the possible occurrence of hemolysis as a factor that contributes to the regulation of erythropoietic homeostasis (Karabulut and Arcagok 2020), we planned to assess endogenous carbon monoxide concentration since it is a more modern and sensitive biomarker for the breakdown of red blood cells (Shahin et al. 2020; Abbott 2022), but due to the limited number of studies addressing this outcome, it was not possible to conduct a meta-analysis. For this reason, we relied on more traditional but widely available markers such as total bilirubin as well as indirect (unconjugated) bilirubin. The pooled analysis showed no statistically significant changes in either marker with exposure to simulated microgravity. Although these findings do not provide strong evidence to support hemolysis, they do not provide sufficient evidence to confidently rule it out either.

The analysis of haptoglobin, another marker of hemolysis (Alayash et al. 2013), also showed no significant change during simulated microgravity exposure. This adds more complexity in interpreting hemolytic activity in this situation, especially in the consideration of whether any hemolysis occurring is primarily intravascular or extravascular in nature. In addition, it should be noted that haptoglobin is an acute phase reactant, which means it is liable to changes in inflammatory conditions like those that may be encountered in spaceflight (Davis et al. 2021; Manis et al. 2022; Tocci et al. 2024). This form of systemic inflammation could increase haptoglobin levels, masking the decreases associated with hemolysis. These aspects limit its reliability as a hemolysis marker in this context.

Several studies that include human subjects exposed to microgravity during spaceflights have reported results that align with the current analysis, like the reductions in hemoglobin concentration, hematocrit levels, and red blood cell mass (RBCM), which have all been documented and are consistent with the results of our study (Kunz et al. 2017; Trudel et al. 2020). While the occurrence of preferential hemolysis of younger erythrocytes after descent from altitude is currently debated (Klein et al. 2021). The agreement between spaceflight data and ground-based analogues increases the confidence in the observed patterns and their applicability in different conditions.

Studies conducted during spaceflight missions have described an early and persistent suppression of erythropoietin (EPO) during microgravity exposure (Leach et al. 1988). This initial decline indicates disruption of the erythropoietic response in advance of any detectable decreases in red blood cell mass. The reduction in EPO concentration has been demonstrated to inversely correlate with central venous pressure, which is a relationship seen in residents of high altitude too, indicating a shared physiological mechanism (Garrett-Bakelman et al. 2019). The severity of spaceflight-associated anemia and recovery time are linked to mission duration, showing the impact of extended erythropoietic suppression, as indicated by the available epidemiological data (Kunz et al. 2017; Trudel et al. 2020).

New papers of less than five years have supported the idea that hemolysis occurs during exposure to microgravity, and they are supported by using recently developed methods to precisely quantify carbon monoxide (CO) elimination as an indicator. One study on actual microgravity showed sustained increases in alveolar CO elimination over the duration of a six-month space mission in 14 astronauts (Trudel et al., 2022). In addition, another study based on ground simulation of microgravity using head-down tilt bed rest also reported increases in CO elimination and bilirubin levels (Culliton et al. 2021), which support the assumption of the occurrence of hemolysis in microgravity, whether in spaceflights or in the simulation models on earth. In contrast, a study which used 21 days of head-down tilt (HDT) bed rest used the same indicator showed no significant increase in CO levels or other hemolytic markers challenging the assumption that hemolysis is a mechanism for red cell loss under simulated microgravity conditions (Trudel et al., 2017).

These contradictory results show the need for more studies to evaluate the nature and magnitude of hemolysis during exposure to microgravity, especially because increased CO levels from space hemolysis can have second-messenger modulatory effects on intracellular processes of the cardiac, vascular, ocular, bone, nutrition, circadian cycle, orthostatic hypotension, brain and muscle systems of astronauts (Afshinnekoo et al. 2020). The use of sensitive and non-invasive indicators as CO elimination should be expanded to detect minor changes in hemolysis that might be missed by traditional hematological parameters (Levitt and Levitt 2015).

It is also important to discuss that the lack of significant changes in the MCV, MCH, and MCHC does not mean that there are no changes in the structure of the red blood cell or its function. Microgravity has been linked with a number of changes in the cytosol of erythrocytes, such as diminished total antioxidant capacity, increased levels of reactive oxygen species (ROS), and lower glutathione concentration, which threaten cell integrity and may shorten the mature RBC lifespan (da Silveira et al. 2020). Moreover, elevated concentrations of phosphatidylcholine and nitric oxide have been reported, which may impair RBC-mediated vasoregulation as well as ATP release and may also contribute to alterations in membrane deformability (Bennett-Guerrero et al. 2007). Research conducted using blood samples from spaceflights indicates that an increase in phosphatidylcholine levels may lead to increased rigidity of the membrane of red blood cells, which is known to limit the functional flexibility of the cells (Ivanova et al. 2006).

These findings show that space medicine still has much potential to advance as current methods of maintaining health in space like Artificial Gravity training sill cannot prevent anemia or other side effects like muscle atrophy (Tran et al. 2021), cardiorespiratory and vascular system alterations (Hoffmann et al. 2021; Möstl et al. 2021; Kramer et al. 2021), and does not affect the neuromuscular secretome (Ganse et al. 2021)., so we can confirm that every physiological change caused by microgravity leads to important clinical effects. Even plasma volume loss, which could be underestimated, has been shown to cause orthostatic intolerance (OI) in astronauts, especially after short-duration spaceflights (Baran et al. 2021). This shows the need for the development of integrated solutions that should ensure fluid replacement, erythropoiesis stimulation, and erythrocyte destruction prevention. Enhancing these countermeasures may contribute not only to the safety of long-duration space missions but also help in the development of treatments for similar conditions on Earth.

This study has multiple strengths, such as the systematic method used to collect and analyze data from several microgravity analog studies. It also provided a detailed and comprehensive overview of the hematological response to simulated microgravity. It included appropriate consideration of multiple outcomes across different domains like blood volume status, erythropoietic activity, red cell morphology, iron metabolism, and hemolysis. This broad coverage improves the explanation of the results and enables a more coherent understanding of space anemia.

Also, there are some limitations that need to be acknowledged. The total number of studies that were included in our analysis was small because of the limited number of studies in this field. Also, the sample size in each study was relatively small, which might decrease their statistical power. In addition, the follow-up periods after reambulation were relatively short for many studies, which limited the ability to give a broader picture for some outcomes.

As for our recommendations for future studies, further research is required to better evaluate the nature and extent of hemolysis during exposure to microgravity. At the same time, there is a need for well-designed studies to develop specific countermeasures intended to maintain red blood cell mass and function. Such measures might include protocols for artificial gravity, exercise programs, pharmacological interventions, or any other methods designed to help the human body adjust to this extreme environment.

### Conclusion

This research provides a comprehensive overview of the hematological changes during simulated microgravity exposure. The findings support the existence of hemoconcentration, erythropoietin suppression, and altered iron metabolism. Many of these changes may be considered a coordinated physiological adaptation rather than a pathological process. However, the possible contributions of low-grade hemolysis require further investigation. Further investigation is also necessary to understand these mechanisms and develop strategies that can aid humans during spaceflights and in relevant clinical situations on Earth.

## Statements and Declarations

### Author contributions

Ahmed Ayman Mohamed Abolyazed led the team, formulated the search strategy, and participated in the literature search, screening, data extraction, risk of bias assessment, and data analysis. He also wrote the manuscript. Yousef Hamdy ElSheeta participated in literature search and screening. Wafaa Ahmed Elbehery participated in data extraction and data analysis. Abdalla Yasser Elnemr participated in risk of bias assessment. Hanan Mahmoud Elsayed supervised all the steps, solved any conflicts and performed a peer review.

### Availability of data and material

All data generated or analyzed during this study are included in this published article and its supplementary information files.

## Acknowledgments

Access to scientific databases was provided through the Egyptian Knowledge Bank (EKB).

## Code availability

**N/A.**

## Ethics approval

**N/A.**

## Consent to participate

**N/A.**

## Consent for publication

**N/A.**

