## supplementary material file for "Effects of Simulated Microgravity on Human Hematological and Anemia-Related Biomarkers: A Systematic Review and Meta-Analysis"

Search Strategy:

(("microgravity" OR "zero gravity" OR "weightlessness" OR "hypogravity" OR "reduced gravity" OR "near-zero gravity" OR "gravitational unloading" OR "simulated microgravity" OR "space flight" OR "spaceflight" OR "space travel" OR "extraterrestrial environment" OR "spacecraft" OR "astronauts" OR "dry immersion" OR "head-down tilt" OR "head down bed rest" OR "parabolic flight")

AND

("anemia" OR "anaemia" OR "iron deficiency" OR "iron deficiency anemia" OR "low hemoglobin" OR "low haemoglobin" OR "hemoglobin" OR "hematocrit" OR "red blood cells" OR "red blood cell count" OR "mean corpuscular volume" OR "MCV" OR "MCH" OR "serum iron" OR "TIBC" OR "total iron-binding capacity" OR "reticulocyte count" OR "hemoglobinuria" OR "aplastic anemia" OR "hemolytic anemia" OR "hypochromic anemia" OR "macrocytic anemia" OR "normocytic anemia" OR "microcytic anemia"))

**Supplementary Fig. 1 Forest plot of changes in Mean Corpuscular Volume during simulated microgravity exposure**

Statistics for each study

| | Difference in means | Standard error | Variance | Lower limit | Upper limit | Z-Value | p-Value |
|---|---|---|---|---|---|---|---|
| Horeau2022 | -1.390 | 0.855 | 0.731 | -3.066 | 0.286 | -1.625 | 0.104 |
| Horeau2024 | 0.000 | 0.541 | 0.293 | -1.060 | 1.060 | 0.000 | 1.000 |
| Lampe1992 | -0.400 | 0.976 | 0.952 | -2.312 | 1.512 | -0.410 | 0.682 |
| Trudel2017 | 0.800 | 0.466 | 0.217 | -0.113 | 1.713 | 1.717 | 0.086 |
| | 0.131 | 0.309 | 0.096 | -0.476 | 0.737 | 0.423 | 0.673 |

Difference in means and 95% CI

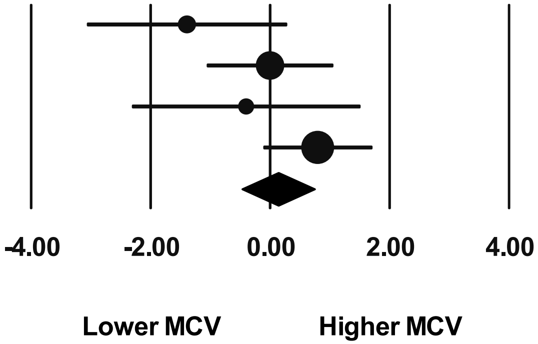


Fixed-effects model used. Heterogeneity: $I^2 = 46.24\%$, $p = 0.134$

**Supplementary Fig. 2** **Forest plot of changes in Mean Corpuscular Hemoglobin during simulated microgravity exposure**

| Study name | Difference in means | Standard error | Variance | Lower limit | Upper limit | Z-Value | p-Value |
|---|---|---|---|---|---|---|---|
| Horeau2022 | 0.010 | 0.327 | 0.107 | -0.631 | 0.651 | 0.031 | 0.976 |
| Horeau2024 | 0.050 | 0.177 | 0.031 | -0.297 | 0.397 | 0.283 | 0.777 |
| Lampe1992 | 0.300 | 0.560 | 0.313 | -0.797 | 1.397 | 0.536 | 0.592 |
| Trudel2017 | 0.100 | 0.164 | 0.027 | -0.222 | 0.422 | 0.609 | 0.543 |
| | 0.078 | 0.111 | 0.012 | -0.139 | 0.295 | 0.704 | 0.482 |

Difference in means and 95% CI

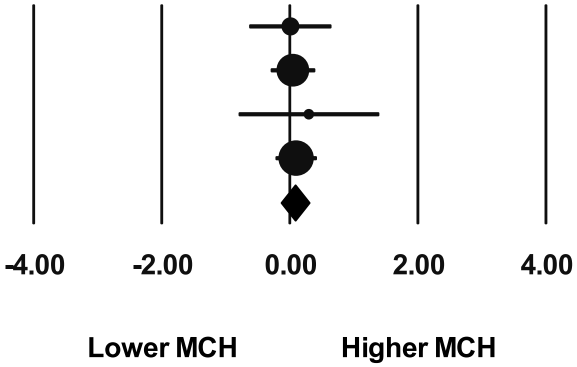


Fixed-effects model used. Heterogeneity: $I^2 = 0.00\%$, $p = 0.970$

**Supplementary Fig. 3** **Forest plot of changes in Mean Corpuscular Hemoglobin Concentration during simulated microgravity exposure**

| Study name | Difference in means | Standard error | Variance | Lower limit | Upper limit | Z-Value | p-Value |
|---|---|---|---|---|---|---|---|
| Horeau2022 | 0.580 | 0.132 | 0.018 | 0.320 | 0.840 | 4.380 | 0.000 |
| Horeau2024 | 0.000 | 0.072 | 0.005 | -0.142 | 0.142 | 0.000 | 1.000 |
| Lampe1992 | 0.800 | 0.413 | 0.170 | -0.009 | 1.609 | 1.938 | 0.053 |
| Ryan2016 | -0.100 | 0.594 | 0.353 | -1.264 | 1.064 | -0.168 | 0.866 |
| Trudel2017 | 0.100 | 0.095 | 0.009 | -0.086 | 0.286 | 1.054 | 0.292 |
| | 0.246 | 0.143 | 0.020 | -0.033 | 0.525 | 1.726 | 0.084 |

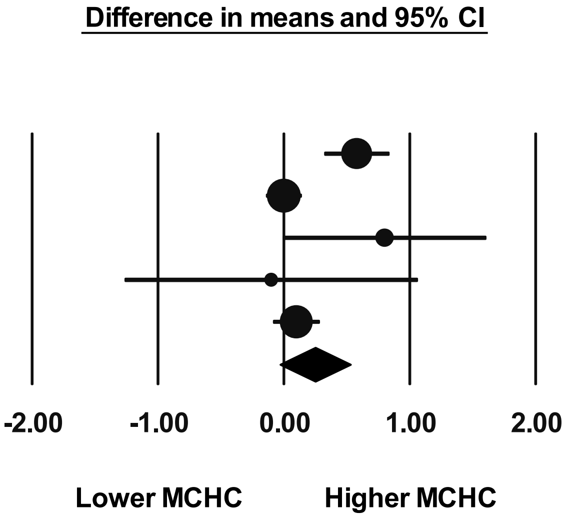


Random-effects model used. Heterogeneity: $I^2 = 77.37\%$, $p = 0.001$

**Supplementary Fig. 4 Forest plot of changes in haptoglobin Concentration during simulated microgravity exposure**

<u>Statistics for each study</u>

| | Difference in means | Standard error | Variance | Lower limit | Upper limit | Z-Value | p-Value |
|---|---|---|---|---|---|---|---|
| Culliton2021 | 0.010 | 0.068 | 0.005 | -0.124 | 0.144 | 0.146 | 0.884 |
| Horeau2022 | 0.100 | 0.077 | 0.006 | -0.051 | 0.251 | 1.296 | 0.195 |
| Nay2020 | 0.100 | 0.062 | 0.004 | -0.021 | 0.221 | 1.624 | 0.104 |
| Ryan2016 | 0.140 | 0.114 | 0.013 | -0.084 | 0.364 | 1.227 | 0.220 |
| Trudel2017 | -0.100 | 0.125 | 0.016 | -0.344 | 0.144 | -0.803 | 0.422 |
| | 0.063 | 0.036 | 0.001 | -0.007 | 0.133 | 1.769 | 0.077 |

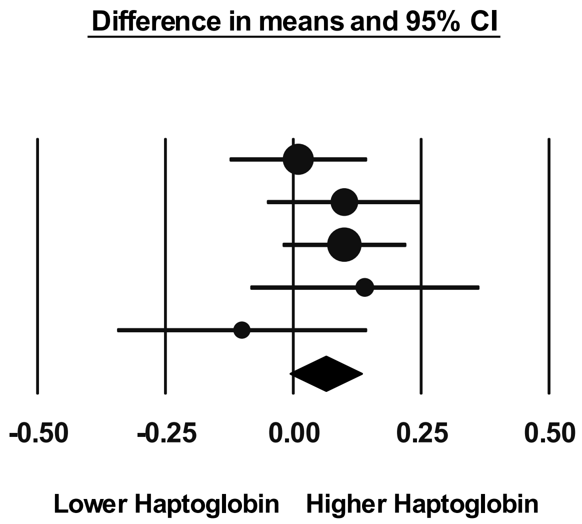


**Fixed-effects model used. Heterogeneity: $I^2$ = 0.00%, p = 0.500**
