## Supplementary figures and images for "Effects of Simulated Microgravity on Human Hematological and Anemia-Related Biomarkers: A Systematic Review and Meta-Analysis"

### Supplementary Fig. 1

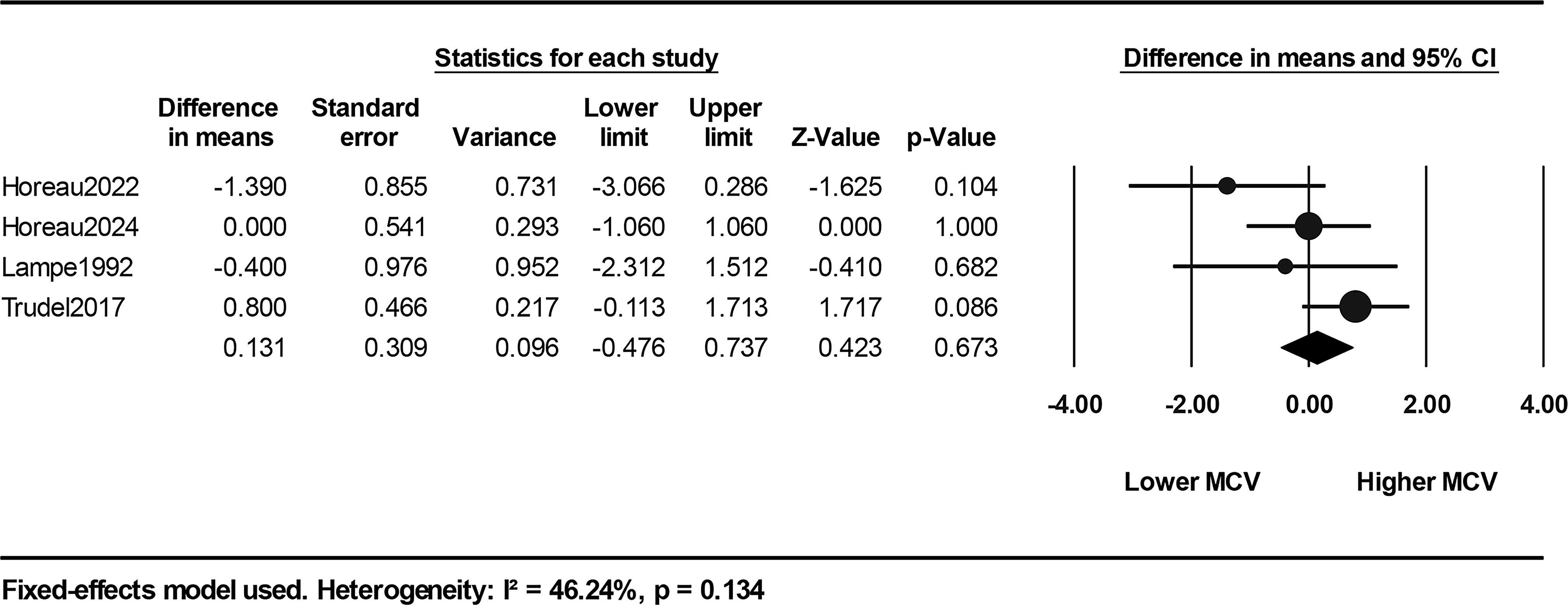

### Supplementary Fig. 2

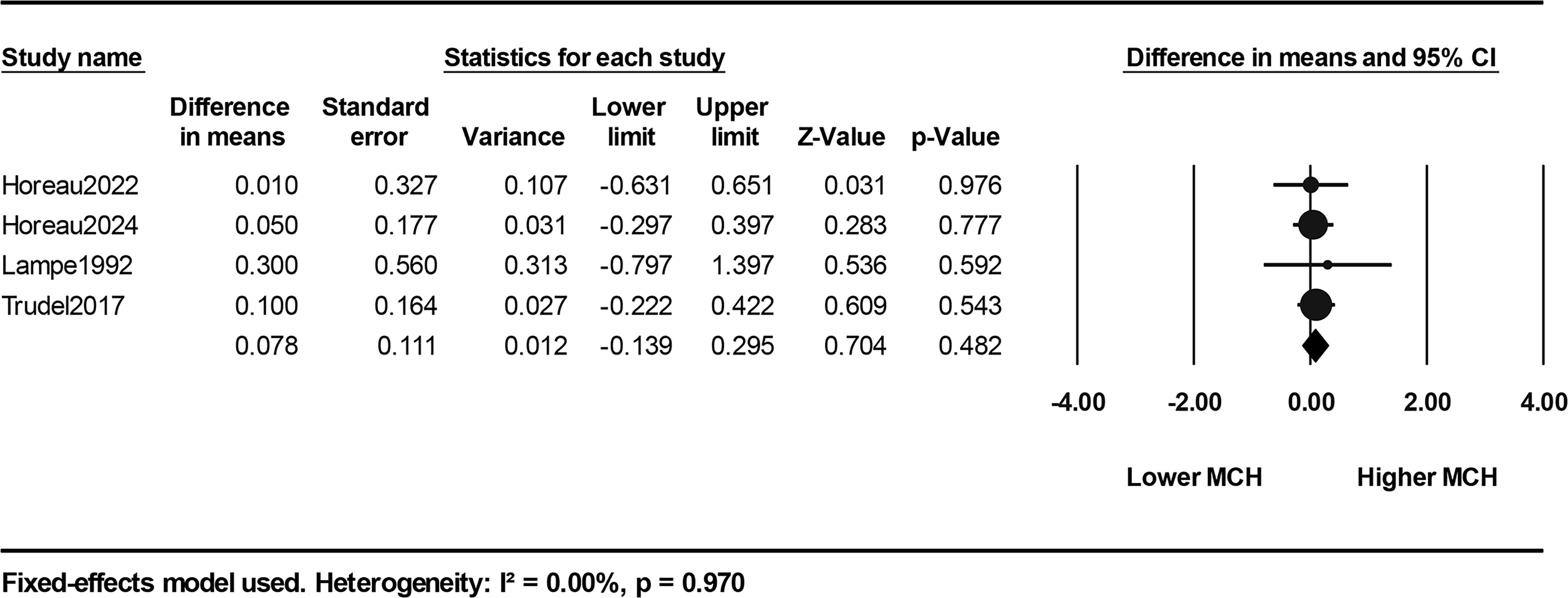

### Supplementary Fig. 3

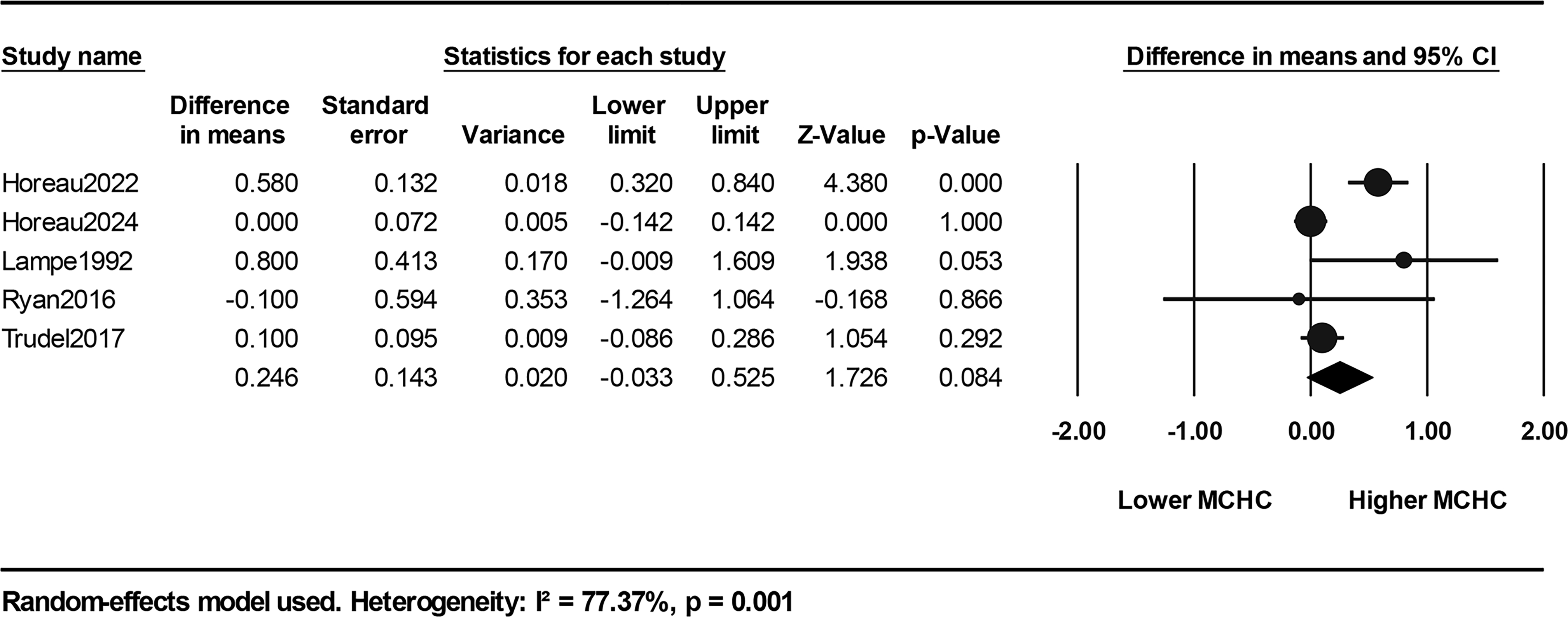

### Supplementary Fig. 4

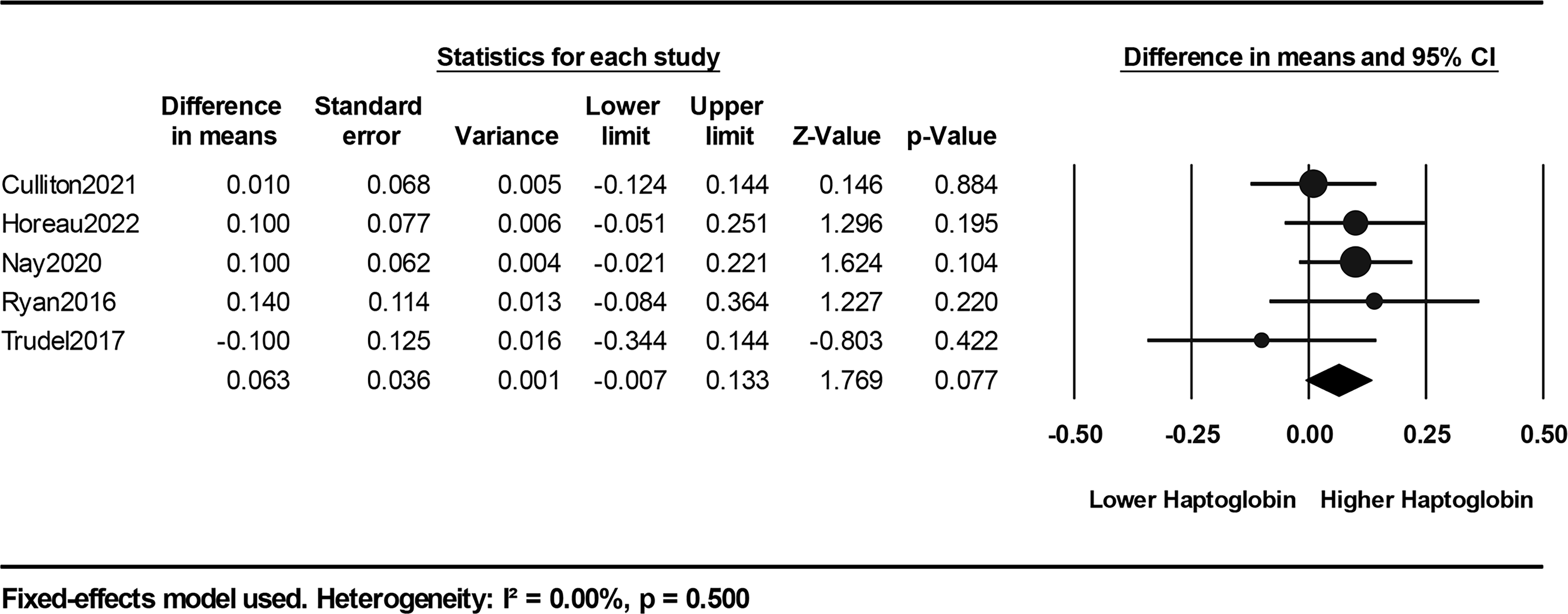
